# Respiration and growth of *Paracoccus denitrificans* R-1 with nitrous oxide as an electron acceptor

**DOI:** 10.1101/2023.10.20.563327

**Authors:** Jiaxian Zhou, Wenfang Deng, Jiapeng Wu, Hua Xiang, Jih-Gaw Lin, Yiguo Hong

**Affiliations:** Institute of Environmental Research at Greater Bay Area; Key Laboratory for Water Quality and Conservation of the Pearl River Delta, Ministry of Education, Guangzhou University, Guangzhou, 510006, China; School of Environmental Science and Engineering, Guangzhou University, Guangzhou, 510006, China; Institute of Environmental Engineering, National Yangming Chiao Tung University, 1001 University Road, Hsinchu City, 30010, Taiwan

**Keywords:** Denitrifier, N_2_O-reduction, electron donor, *nos*Z, *Paracoccus denitrificans* R-1

## Abstract

In the nitrogen biogeochemical cycle, the reduction of nitrous oxide (N_2_O) to N_2_ by N_2_O reductase, which is encoded by *nos* gene cluster, is the only biological pathway for N_2_O consumption. However, the capability and mechanisms of microbial N_2_O reduction are poorly understood. In this study, we investigated the ability to obtain energy for growth of *Paracoccus denitrificans* R-1 by coupling the oxidation of various electron donors to N_2_O reduction. This strain has strong N_2_O reduction capability, and the average N_2_O reduction rate was 5.10±0.11×10^-9^ μmol·h^-1^·cell^-1^ under anaerobic condition at 30℃ using acetate as the electron donor in a defined medium. This reduction was accompanied by the stoichiometric consumption of acetate over time when N_2_O served as the sole electron acceptor and the reduction can yield energy to support microbial growth, suggesting that microbial N_2_O reduction is an electron transport process. Cu^2+^, silver nanoparticles, O_2_, and acidic conditions can strongly inhibit the reduction, whereas NO_3_^-^ or NH_4_^+^ can promote it. Genomic analysis showed that the gene cluster encoding N_2_O reductase of *P. denitrificans* R-1 was composed of *nos*R, *nos*Z, *nos*D, *nos*F, *nos*Y, and *nos*L, and *nos*Z, which was identified as clade I. The respiratory inhibitors test indicated that the pathway of electron transport for N_2_O reduction was different from that of the traditional electron transport chain for aerobic respiration. These findings suggest that modular N_2_O reduction by *P. denitrificans* R-1 is linked to the electron transport chain and energy conservation, and that dissimilatory N_2_O reduction is a form of microbial anaerobic respiration.

**IMPOETANCE:** In the nitrogen biogeochemical cycle, the reduction of N_2_O to N_2_ by N_2_O reductase, which is encoded by *nos genes*, is the only biological pathway for N_2_O consumption. However, the capacity and mechanisms of microbial N_2_O reduction are poorly understood. We investigated the ability to obtain energy for growth of Paracoccus denitrificans R-1 by coupling the oxidation of various electron donors to N_2_O reduction. Our study showed that the nosZ type I bacterium, P. denitrificans R-1, can respire N_2_O as the sole electron donor. Thus, the modular N_2_O reduction process of clade I denitrifiers not only can consume N_2_O produced by themselves but can also consume the external N_2_O generated from non-denitrification biological or abiotic pathways under suitable conditions, which is critical for controlling the release of N_2_O from ecosystems into the atmosphere.

## INTRODUCTION

Nitrous oxide (N_2_O), a colorless, stable gas, has an average lifetime of 120 years in the atmosphere and is ultimately decomposed by ultraviolet light (1). As one of the most important forms of nitrogen pollution, N_2_O is currently the third largest greenhouse gas (GHG) emitted and the largest anthropogenic stratospheric ozone-depleting substance (Davidson and Kanter, 2014). Although N_2_O only accounts for approximately 0.03% of total GHG emissions, it has a nearly 300-fold greater potential for global warming based on its radiative capacity compared to that of carbon dioxide (CO_2_) (2). Therefore, controlling N_2_O emissions is essential for curbing global warming and climate change.

N_2_O emissions from soil involve a variety of biological pathways, and it has been estimated that more than 65% of atmospheric N_2_O is derived from microbial N transformations, mainly through the processes of nitrification and denitrification (3). Among them, denitrification is generally considered the largest source of N_2_O, and depending on the types of microorganisms involved and environmental conditions, this process can serve not only as a source of N_2_O but also as a sink for N_2_O (Thomson et al., 2012). Denitrification is the respiratory reduction of nitrogen oxides (NOx) and enables the survival and reproduction of facultative aerobic bacteria under oxygen-limiting conditions. In this process, nitrate (NO_3_ ^-^) is converted into molecular nitrogen (N_2_) via nitrite (NO_2_^-^) and the gaseous intermediates, nitric oxide (NO) and nitrous oxide (N_2_O) (4).

In contrast to the large number of N_2_O production pathways and enzymes, only one enzyme is involved in biological N_2_O consumption. This Cu-dependent enzyme is known as N_2_O reductase (*nosZ*) (2). In typical denitrifying microorganisms (such as *Proteobacteria* of α-, β- and γ-classes), NosZ has long been considered the only enzyme that can reduce N_2_O to N_2_, which is called “clade Ⅰ NosZ”. However, an unprecedented *nos* gene cluster with a novel *nosZ* containing an additional c-type heme domain at the C terminus was discovered, which was called “clade Ⅱ NosZ,” and, which has been identified in a broad range of microbial taxa extending beyond bacteria to archaea (5, 6).

According to the current study, clade Ⅰ NosZ and clade Ⅱ NosZ are two different phylogenetic groups of the NosZ protein. Additionally, the types of clade Ⅱ NosZ microorganisms are more complex compared to clade Ⅰ NosZ microorganisms, and clade II NosZ contains some genes that are not present in clade I NosZ organisms (6, 7). The clade Ⅰ *nos* gene cluster contains NosZ, -R, -X, -C, -D, -F, -Y, and, -L, whereas the clade Ⅱ *nos* gene cluster contains NosB, -Z, -G, -H, -C, -C1, -C2, -D, -F, -Y, and -L; NosB, -C1, -C2, and, -G in clade II replace the missing NosR in the electron transport chain (2). Recently, an increasing number of NosZ-containing microorganisms have been reported to grow via anaerobic N_2_O respiration, with N_2_O as the only electron acceptor, including *Bacillus vireti* (8), *Enifer meliloti* 1021 (9), *Azospira* sp. strain I13 (10), and *Gemmatimonas aurantiaca* strain T-27 (11).

Microorganisms containing *nosZ* are abundant in the environment. However, only a few N_2_O-reducing bacteria have been successfully isolated and identified, and there is a limited understanding of their reducing mechanisms. In this study, *Paracoccus denitrificans* R-1, a denitrifying bacterium isolated from the Xinfeng Sewage Plant in Taiwan, was used to explore its N_2_O reduction ability. Our results showed that the reduction of N_2_O by *P. denitrificans* R-1 is a new pathway for N_2_O respiration.

## Materials and methods

### Media, strain, and cultivation

*P. denitrificans* R-1 was isolated from sludge of the Taiwan Xifeng Sewage Treatment Plant and stored at the Guangdong Provincial Microbial Strain Preservation Center (GDMCC 1.2910). This strain was stored at -80°C and was pre-cultured aerobically in nutrient medium (pH 7.0) containing 10 g L^-1^ NaCl, 5 g L^-1^ Bacto Peptone, and 5 g L^-1^ Oxoid™ Lab-Lemco meat extract at 30°C with shaking at 150 rpm. Then, when the culture reached the exponential phase, it was inoculated into vials of N-free denitrifying medium (N-free DM), which contained 10 g L^-1^ of Na_2_HPO_4_·12H_2_O, 1.5 g L^-1^ of KH_2_PO_4_, 0.1 g L^-1^ of MgSO_4_·7 H_2_O, 4.7 g L^-1^ of sodium acetate, and 2 mL L^-1^ of a trace metal solution (12). The cell densities (calibrated with the absorbance value of OD_600_) of the culture were measured at 600 nm using a spectrophotometer (Shimadzu Enterprise Management Co., LTD).

### Incubation

Aerobically grown cells in DM were harvested by centrifugation, washed twice, and resuspended in fresh N-free DM (initial pH 7.5). Then, different organic substances were supplied as electron donors, after which the cells were dispensed into 60-mL serum bottles (30 mL per bottle) and ensure that the initial OD_600_ of the bacteria in the serum bottles are 0.05, crimp-sealed with rubber septa and aluminum caps to ensure an airtight system. The headspace of the serum vials (30 mL volume) was subsequently replaced with 10% N_2_O (He: N_2_O = 9:1) to analyze the N_2_O reduction capability of strain R-1. Different electron donors (carbon sources) were used to explore the coupling of the oxidation of electron donors (carbon sources) with the reduction of electron acceptors (N_2_O). Different respiratory inhibitors were added to explore possible electron transport pathways involved in N_2_O reduction. Furthermore, the effects of temperature, pH, dissolved oxygen, heavy metal ions, and nitrogen substrates (NO_3_ ^-^ and NH_4_ ^+^) on the reduction of N_2_O were conducted.

### N_2_O measurement

The N_2_O amount in the incubation bottle was composed of two parts, one in the headspace and the other dissolved in liquid medium. Three parallel incubations were performed for each sample. After incubation, 50% ZncL_2_ was used to inactivate the bacterial cells. The N_2_O concentration was measured by manually injecting 3–4 mL diluted headspace gas into a gas chromatograph equipped with an HP-PLOT/Q column and an electron capture detector (GC-2014C, Shimadzu Enterprise Management Co., LTD, China). Headspace N_2_O concentration (C_G,_ μmol/L) in serum bottle is calculated by the following equation 1:

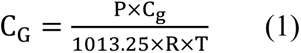

Where P is the atmospheric pressure in the serum bottle, C_g_ (ppm) (1ppm=1μmol·mol^-1^) is the concentration of N_2_O in the headspace measured with GC and R is the ideal gas constant, i.e., 0.082057 Latm·(mol·K)^-1^. T (K) is the temperature of the water sample at headspace equilibrium.

The N_2_O concentration dissolved in the serum bottle liquid (C_L_, μmol·L^-1^) is calculated by the following equation 2:

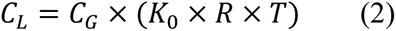

K_0_ (mol·(L·atm) ^−1^) denotes the equilibrium constant which can be calculated by Weiss formula (13).

The total amount of N_2_O (Q, μmol) in the serum bottle is calculated by the following equation 3:

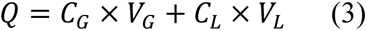

Where V_G_ and V_L_ are the volumes of gas and liquid, respectively.

### Electron donor measurement

Small-molecule organic substances including acetate and lactate were used as electron donors in this study. The cultures were filtered through a 0.22 μm membrane and the filtrate was used to measure the concentration of electron donors. Acetate and lactate were measured using an ICS-1100 series ion chromatograph (Thermo Fisher Scientific, Waltham, MA, USA) equipped with a polysulfonate ion-exclusion column (Metrosep A Supp 5). The eluent contained the following: 3.2 mmol L^-1^ Na_2_CO_3_, 0.8 mM NaHCO_3_, and 3% MeOH. The experiment was performed under the condition of 25°C and 7.3 MPa.

### DNA extraction, sequencing, and genomic analysis

*P. denitrificans* R-1 was inoculated into the nutrient medium and cultured until the logarithmic stage. The bacteria were collected in sterilized centrifuge tubes and stored at -80°C, with a mass of approximate 2 g. Genomic DNA extraction and sequencing of this strain performed by Suzhou GENEWIZ Biotechnology Co., Ltd. After obtaining high-quality genomic DNA, the fragments were randomly broken down into the corresponding length fragments to construct a library. The qualified libraries were sequenced (150-bp paired-end sequencing) on the NovaSeq system. Detailed information regarding genomic sequencing can be found in our previous study (14).

The genome analysis process consisted of four steps: (1) data quality control: preprocessing of the original data obtained by sequencing. Low-quality data were filtered, and splice sequences were removed to prevent low-quality data from having a negative impact on subsequent analyses. Clean data obtained after data preprocessing were used for subsequent analyses. The software used for the quality statistics of the second-generation sequencing data was adapted (v 1.9.1). (2) Genome assembly: HGAP4 software was used to assemble the third-generation sequencing data. After the assembly was completed, the quality control of the second-generation sequencing data was compared with that of the third-generation assembly results, and the final assembly results were obtained using Pilon. (3) Prediction of coding and non-coding genes: after assembly, coding and non-coding RNAs were predicted using the Prodigal software (v3.02). (4) Gene function annotation: the predicted protein sequence of the coding gene was compared with the protein sequences contained in each database. The databases used in this study mainly included Cluster of Orthologous Groups (COG), Gene Ontology (GO), Carbohydrate-Active Enzymes Database (CAZy), and Kyoto Encyclopedia of Genes and Genomes (KEGG).

## RESULTS

### N_2_O reduction coupled to oxidation of electron donors

To explore the relationship between N_2_O reduction and electron donor oxidation, changes in the concentrations of both electron donors and N_2_O in the culture system were analyzed simultaneously. In the culture with sodium acetate as the electron donor, the N_2_O content decreased from the initial 83.81±1.65 μmol to 0.58±0.24 μmol after 16 h of culturing. N_2_O content decreased by 83.23±1.89 μmol. In addition, sodium acetate decreased from 173.45±4.77 μmol to 78.84±12.44 μmol during culture, and sodium acetate decreased by 94.61±17.21 μmol (Fig. 1A). In the culture with sodium lactate as the electron donor, the N_2_O content decreased from the initial 89.63±0.28 μmol to 0.77±0.03 μmol at 16 h, N_2_O content decreased by 88.86±0.31 μmol, and sodium lactate decreased from 105.99±8.28 μmol to 63.13±0.74 μmol in 0-16 h, sodium lactate decreased by 42.86±7.54 μmol (Fig. 1B). According to the fitting equations (R^2^=0.9764 for acetate and R^2^=0.9485 for lactate), there was a significant linear relationship between the reduction of N_2_O and the oxidation of the electron donors of acetate or lactate.

**Fig. 1.**
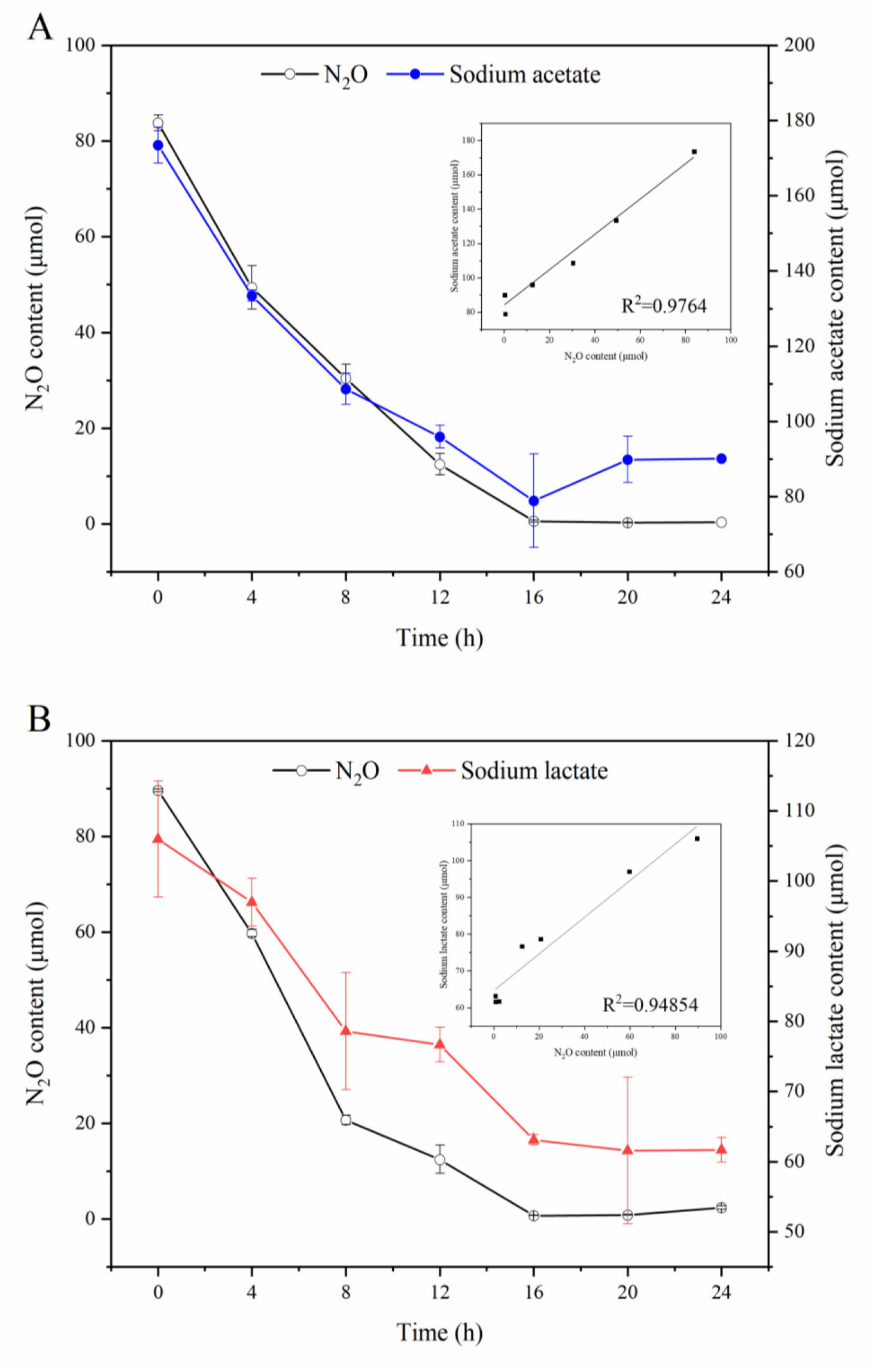
*P. denitrificans* R-1 for N_2_O reduction coupled with electron donor oxidation. Sodium acetate as electron donor (A) and sodium lactate as electron donor (B). Data points are averages of duplicate experiments and error bars represent standard deviations.

### Growth of *P. denitrificans* R-1 with N_2_O as sole electron acceptor

In a culture experiment using sodium acetate as the electron donor and N_2_O as the sole electron acceptor, *P. denitrificans* R-1 exhibited N_2_O consumption and growth (Fig. 2). The consumption of N_2_O was slow from 0–4 h, and the concentration of N_2_O decreased rapidly from 4–20 h. The N_2_O concentration virtually remained unchanged from 20– 24 h. In this process, the N_2_O content decreased from the initial 83.78±2.88 μmol to 5.47±0.72 μmol, and the average N_2_O consumption rate was 5.10±0.11 × 10^-9^ μmol·h^-1^·cell^-1^. However, no significant change in the concentration of N_2_O was observed in the no-cell incubation, indicating that the reduction of N_2_O in the experimental group was not caused by spontaneous degradation or transformation of N_2_O but was consumed by *P. denitrificans* R-1. No N_2_O release was observed during culturing. Furthermore, N_2_O reduction was accompanied by the growth of *P. dienitrificans* R-1 in the culture. The growth was slow during 0–4 h but became relatively faster after 4 h and remained stable for 20–24h. The OD_600_ value of *P. denitrificans* R-1 increased from 0.0675 to 0.1925 within 24 h (an increase of 0.1858). However, the OD_600_ value virtually did not change within 24 h in the control group without N_2_O, suggesting that the growth of *P. denitrificans* R-1 was due to energy conservation during N_2_O reduction.

**Fig. 2.**
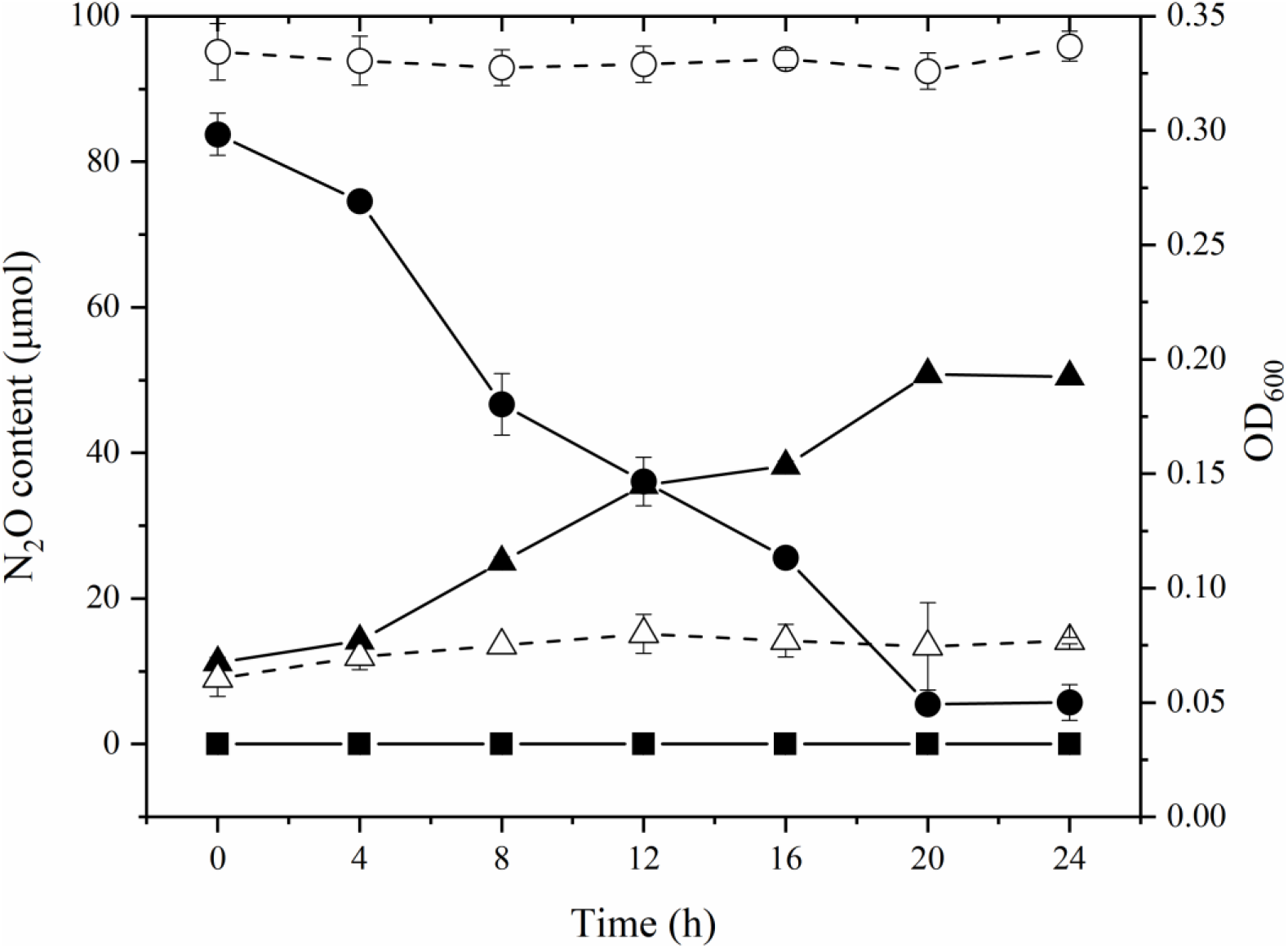
Growth of *P. denitrificans* R-1 during N_2_O reduction. In control vessels without N_2_O, cell numbers did not increase. No N_2_O reduction occurred in control cultures that received no inoculum. ●, N_2_O; ▪, He, no cells; ▴, cells = OD_600_ nm; ○, N_2_O, no cells; △, cells = OD_600_ nm, no N_2_O. Data points are averages of duplicate experiments and error bars represent standard deviations.

### Effect of respiratory inhibitors on N_2_O reduction

To explore the electron transport chain components that are possibly involved in N_2_O reduction by *P. denitrificans* R-1, three respiratory inhibitors (dicoumarin, rotenone, and antimycin A) were used to explore the effect of N_2_O reduction by *P. denitrificans* R-1. Among these, dicoumarin inhibits electron transport from vitamin K to quinones, rotenone blocks electron transport from NADH to CoQ, and antimycin A inhibits electron transport from QH_2_ to cytochrome C_1_ (15). All three inhibitors had no significant effect on the N_2_O reduction process by *P. denitrificans* R-1 (Fig. 3), suggesting that N_2_O reduction by *P. denitrificans* R-1 had different electron transport components from the traditional one involving complexes I and II. Regarding which electron transport components were involved in N_2_O reduction by this strain, further study using new methods is necessary.

**Fig. 3.**
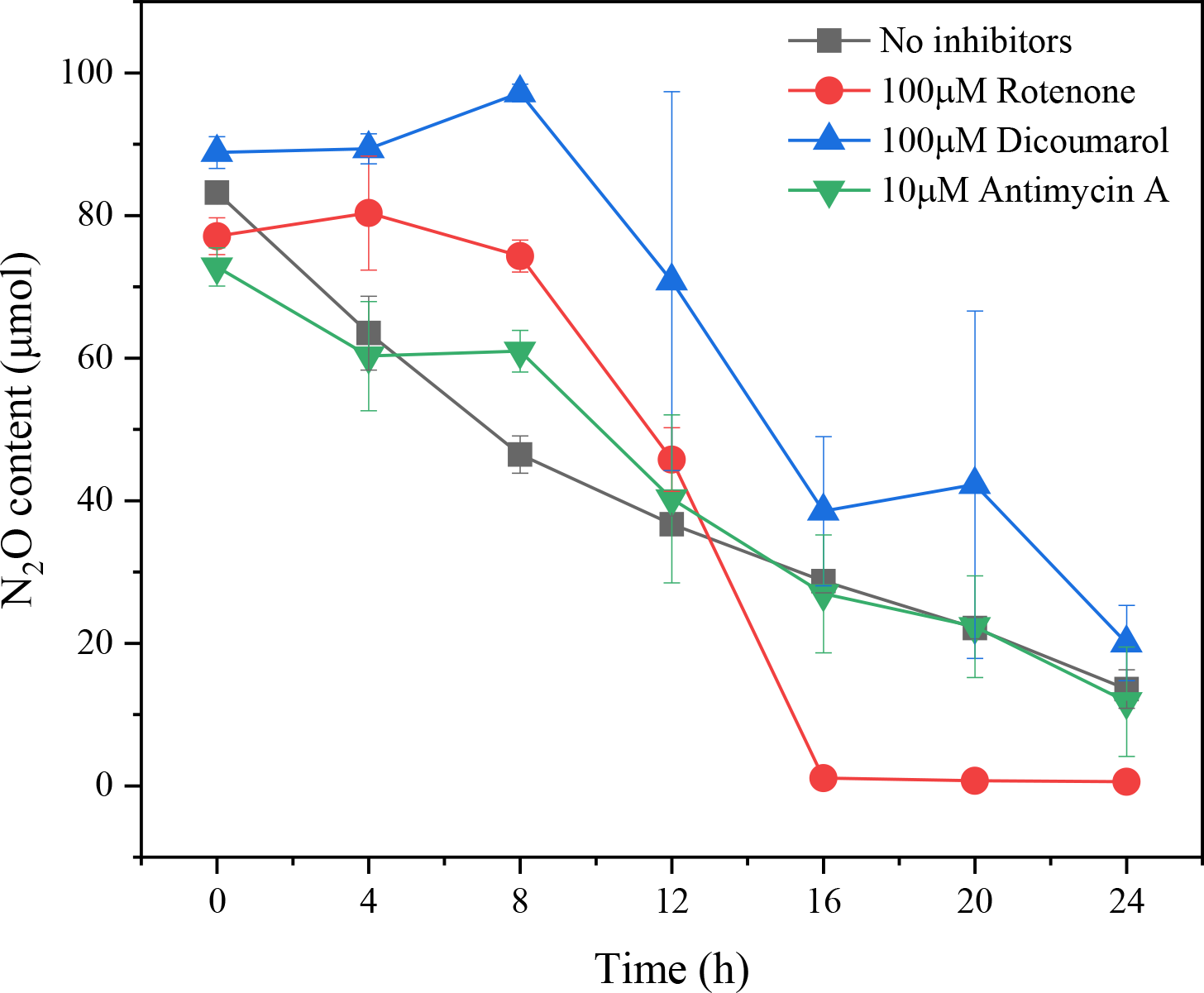
Respiratory inhibition experiment of *P. denitrificans* R-1. Data points are averages of duplicate experiments and error bars represent standard deviations.

### Composition and characteristics of gene cluster of *P. dienitrificans* R-1

A phylogenetic evolutionary tree was constructed using *nosZ* sequences from different bacterial strains collected from the NCBI database (Fig. 4). The evolutionary tree was divided into two clusters, clade I and clade II, with *P. denitrificans* R-1 (bold) distributed in clade I. The differences between clades I and II are not only reflected in the phylogeny of the NosZ protein, but also in the composition and structure of their respective *nos* gene clusters (16). The genomic locus encoding NosZ is a part of the *nos* gene cluster, which also includes genes encoding helper proteins required for the maturation and function of NosZ (6). *P. denitrificans* R-1 *nos* gene cluster consisted of *nosR*, *nosZ*, *nosD*, *nosF*, *nosY*, and *nosL*, which is a common pattern in *nos* gene cluster of clade I microorganisms. However, clade II microorganisms have more complex and diverse *nos* gene clusters than those of clade I microorganisms (Fig. 5). Because *nosD*, -*F*, and -*Y* are considered to be the strongest conserved genes in *nos* gene clusters, they exist in both clade I and clade II type microorganisms (17).

**Fig. 4.**
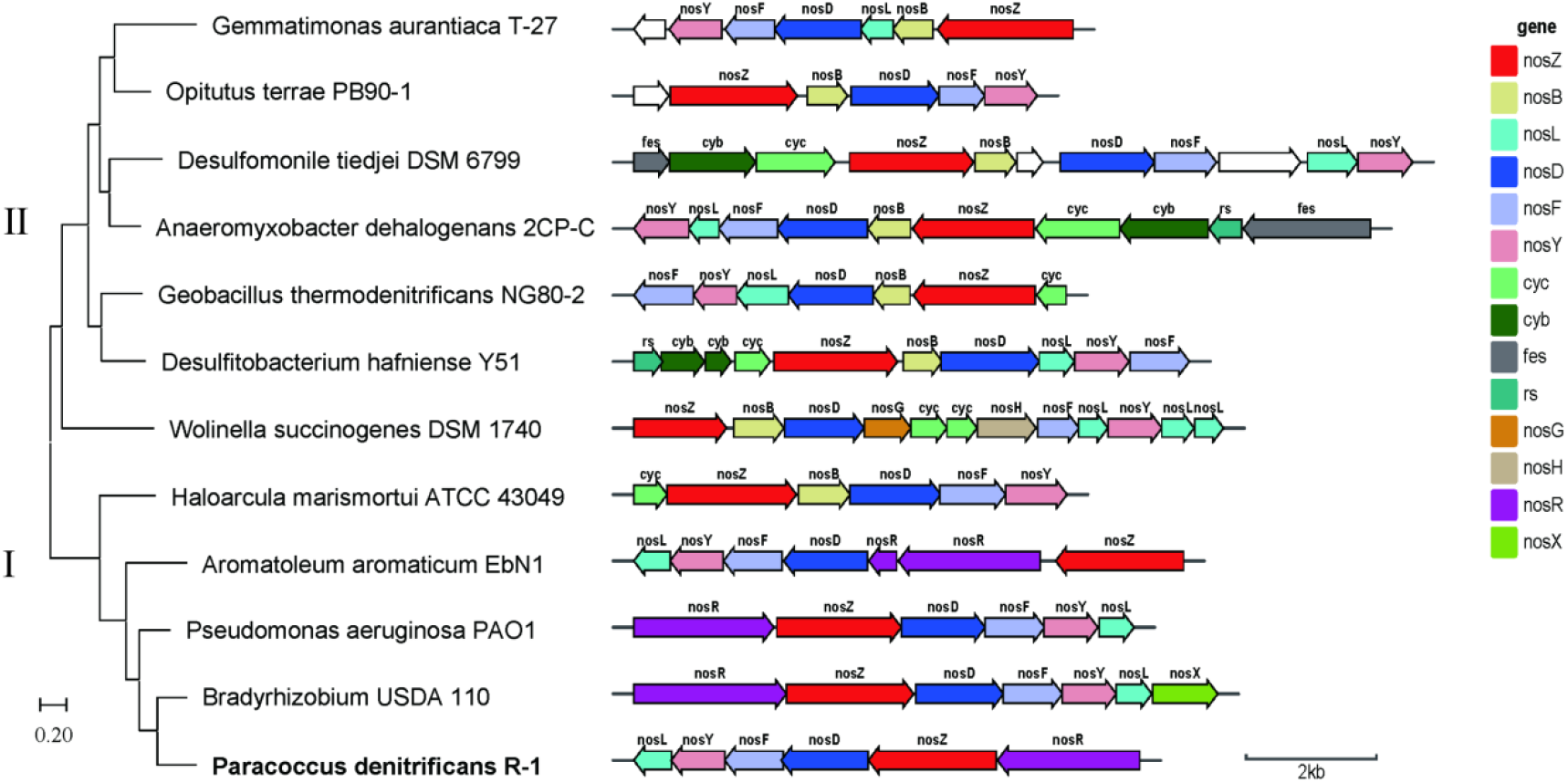
Phylogenetic tree based on NosZ protein sequence and comparison of *nos* gene clusters in different clades of NosZ. For each gene cluster, *nosZ* and its accessory genes (BDFGHLRXY) are labeled and colored according to homology across different gene clusters. Additional proteins, including iron-sulfur-binding proteins (FeS), Rieske iron-sulfur proteins (S), and b- and c-type cytochromes (Cyb and Cyc, respectively) are also labeled. Noncolored genes denote open reading frames with no orthologs in other *nos* gene clusters, and scale bar at lower-right indicates gene size.

**Fig. 5.**
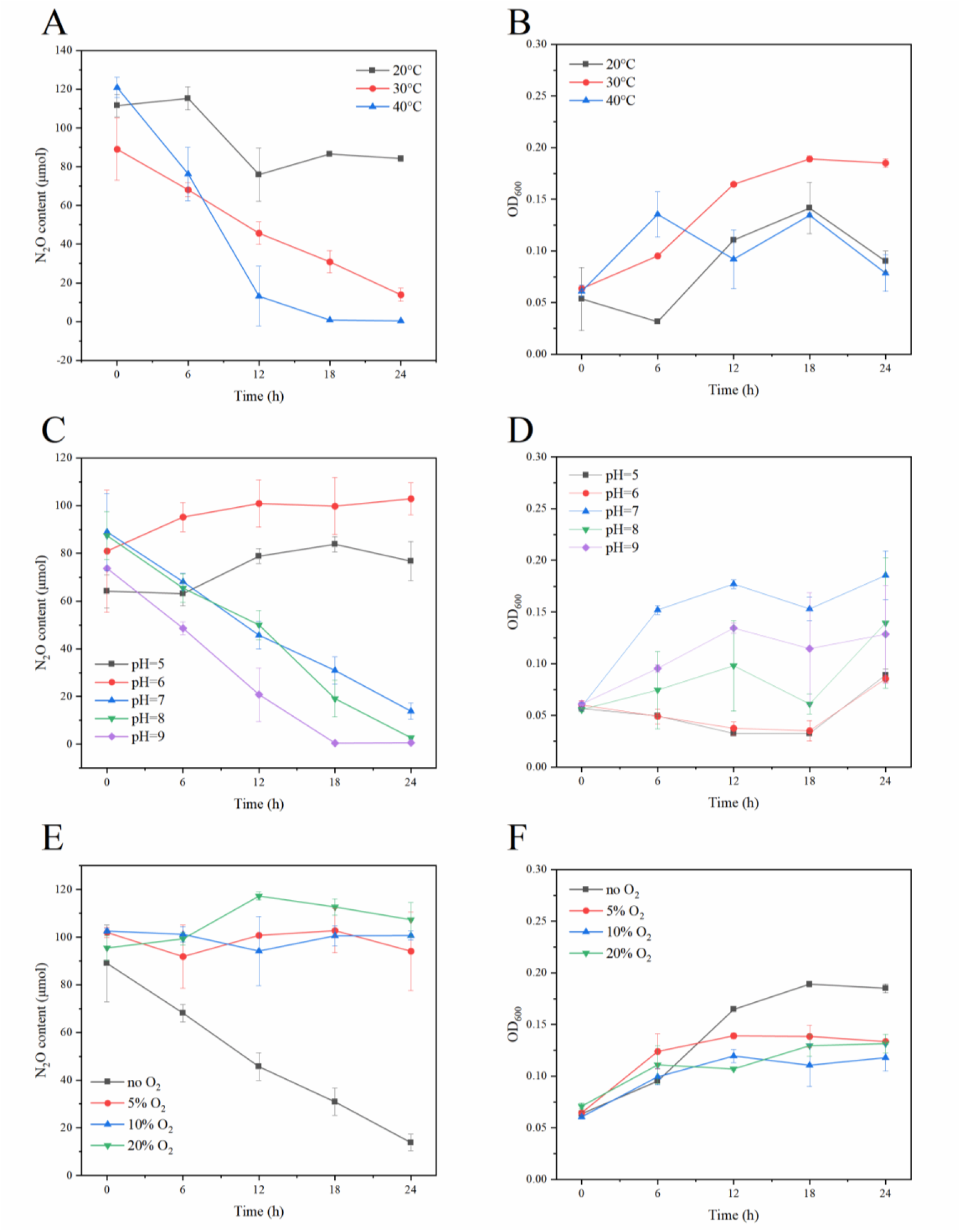
Effects of temperature, pH, and O_2_ on N_2_O reduction and growth of *P. denitrificans* R-1. N_2_O consumption curve (A) and strain growth curve (B) at different temperatures. N_2_O consumption curve (C) and strain growth curve (D) at different pH values. N_2_O consumption curve (E) and strain growth curve (F) under different O_2_ concentrations. Data points are averages of duplicate experiments and error bars represent standard deviations.

### Effects of temperature, pH, and O_2_ on N_2_O reduction

The average N_2_O reduction rate of *P. denitrificans* R-1 was 2.00±0.30×10^-9^ μmol·h^-1^·cell^-1^ (0-24 h), 5.50±0.53×10^-9^ μmol·h^-1^·cell^-1^ (0-24 h), and 1.17±0.30×10^-8^ μmol·h^-1^·cell^-1^ (0-18 h) under the condition of 20, 30 and 40℃, respectively (Fig. 5A). The OD_600_ value for growth of *P. denitrificans* R-1 was 0.037±0.015, 0.129±0.006, and 0.018±0.017 at 20, 30, and 40℃, respectively (Fig. 5B). These results suggest that the N_2_O reduction rate of *P. denitrificans* R-1 was promoted with an increase of temperature in a limited range, but the growth of bacteria was most obvious at 30℃.

At pH 5.0 and 6.0, no N_2_O consumption or cell growth was observed within 24 h, suggesting that the reduction of N_2_O by *P. denitrificans* R-1 was not favorable under acidic conditions (Fig. 5C). At pH 7.0, 8.0, and 9.0, *P. denitrificans* R-1 was able to reduce N_2_O normally (Fig. 5C), indicating that the N_2_O-reducing ability of the strain was activated under neutral or alkaline conditions. Growth was optimal at pH 7.0, but it was inhibited under acidic or alkaline conditions (Fig. 5D).

N_2_O oxidoreductase (NosZ) was thought to be sensitive to O_2_ (18). Three gradient concentrations of O_2_ (5, 10, and 20%) were selected for culturing *P. denitrificans* R-1. No N_2_O consumption was observed after 24 h culture under all three gradient concentrations of O_2_ (Fig. 5E), suggesting that the O_2_ can strongly inhibit the N_2_O reduction of *P. denitrificans* R-1.

### Effects of NO_3_^-^ and NH_4_^+^ on N_2_O reduction

In the culture where different concentrations of NO_3_^-^ were added, the N_2_O reduction rate by *P. denitrificans* R-1 increased (Fig. 6A). The average N_2_O reduction rate by *P. denitrificans* R-1 was 1.53±0.69×10^-8^ μmol·h^-1^·cell^-1^, which was approximately 3.5 times higher than that in the culture without NO_3_^-^ (4.39±0.13×10^-9^ μmol·h^-1^·cell^-1^). This suggests that the N_2_O reduction of *P. denitrificans* R-1 can be promoted by NO_3_^-^ in the culture. Moreover, in the culture with NO_3_^-^ (except for the addition of 20 mg·L^-1^ NO_3_^-^), the growth rate of *P. denitrificans* R-1 was higher than that in the culture without NO_3_^-^(Fig. 6B), possibly because the strain can use NO_3_^-^ as an electron acceptor to obtain energy for growth. Similarly, the N_2_O reduction capability of *P. denitrificans* R-1 improved in cultures with different concentrations of NH_4_^+^ (Fig. 6C). The average N_2_O reduction rate by *P. denitrificans* R-1 was 1.39±0.86 ×10^-8^ μmol·h^-1^·cell^-1^, which was approximately 3 times higher than that in the culture without NH_4_^+^ (4.39±0.13×10^-9^ μmol·h^-1^·cell^-1^). The growth of strains with NH_4_^+^ added was significantly higher than that without NH_4_^+^ (Fig. 6D). Therefore, both NO_3_^-^ and NH_4_^+^ in the culture improved N_2_O reduction by *P. denitrificans* R-1.

**Fig. 6.**
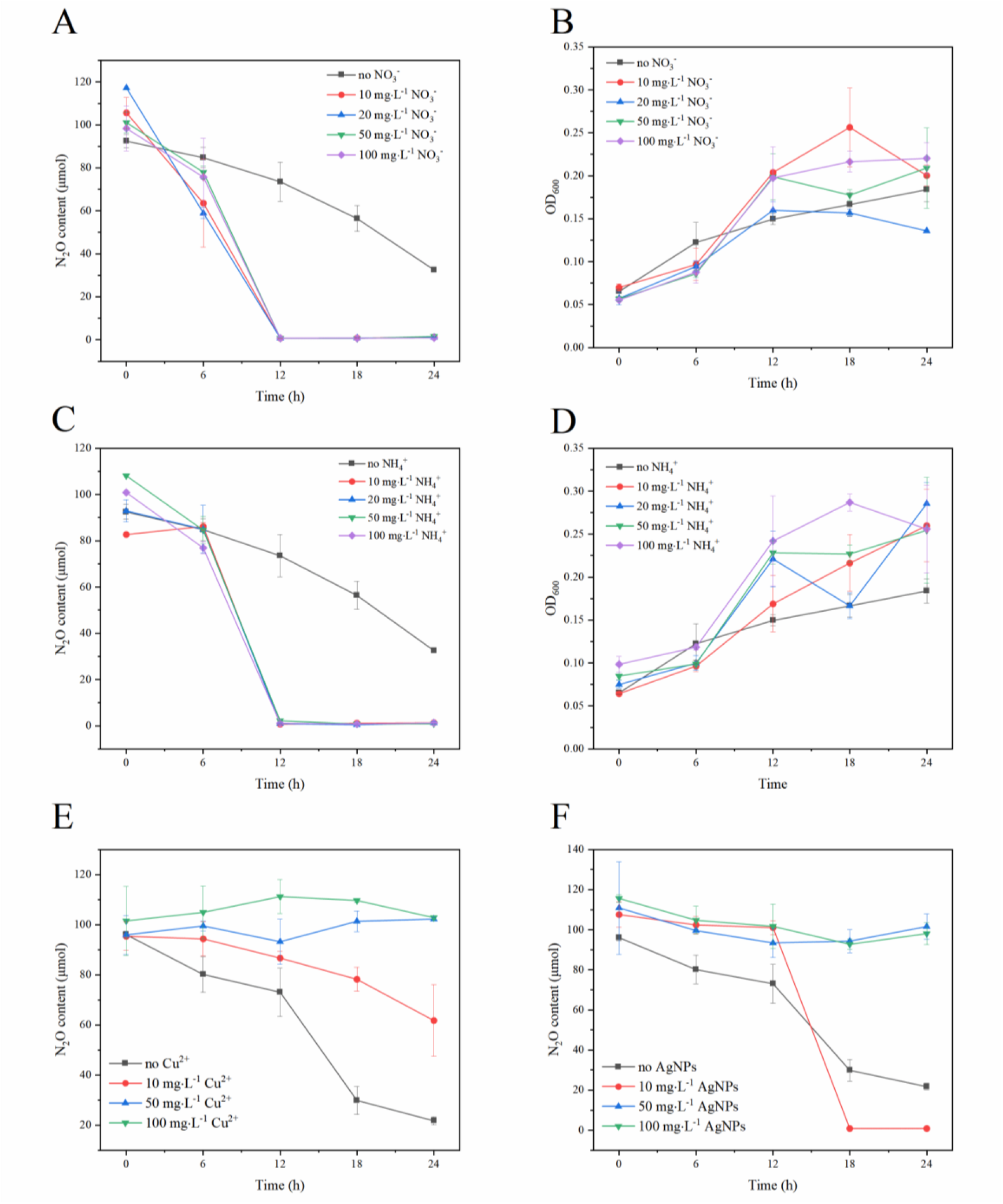
Effects of adding different nitrogen substrates and heavy metal ions on N_2_O reduction and growth of *P. Denitrificans* R-1. After addition of metal ions, turbidity, and color of medium had an impact on OD_600_ determination, so OD_600_ value of growth with addition of heavy metal ions was not measured. N_2_O-reduction process curve (A) and growth curve (B) of *P. denitrificans* R-1 under different concentrations of NO_3_^-^. N_2_O-reduction process curve (C) and growth curve (D) of *P. denitrificans* R-1 under different concentrations of NH_4_^+^. N_2_O-reduction at different Cu+ concentrations (E). N_2_O-reduction at different AgNPs concentrations (F). Data points are averages of duplicate experiments with error bars representing standard deviations.

### Effects of heavy metal ions on N_2_O reduction

Trace amounts of Cu are believed to promote microbial growth and this element is an important component of *NosZ* (2, 19). With the development and application of nanomaterials, silver nanoparticles (AgNPs) have become the most widely used owing to their superior bactericidal abilities (20, 21). Two types of heavy-metal ions, Cu^2+^ and AgNPs, were selected to analyze their effects on N_2_O reduction. When the the concentration of Cu^2+^ was 10 mg·L^-1^, the N_2_O reduction rate decreased to 2.46±0.35×10^-9^ μmol·h^-1^·cell^-1^ compared to the rate of 5.42±0.01 ×10^-9^ μmol·h^-1^·cell^-1^ in the culture without Cu^2+^. When the concentration of Cu^2+^ increased to 50 mg·L^-1^ and 100 mg·L^-1^, the reduction of N_2_O was not completely detected (Fig. 6E). These results demonstrate that Cu^2+^ strongly inhibited N_2_O reduction of *P. denitrificans* R-1. In contrast, the reduction rate of N_2_O increased from 5.42±0.01×10^-9^ μmol·h^-1^·cell^-1^ to 1.04±0.35 ×10^-8^ μmol·h^-1^·cell^-1^ when the concentration of AgNPs was 5 mg·L^-1^ (Fig. 6F), indicating that a low concentration of AgNPs was able to promote N_2_O reduction by *P. denitrificans* R-1. However, N_2_O reduction was greatly reduced when the concentration of AgNPs was increased to 10 or 20 mg·L^-1^, (Fig. 7F), suggesting that the high concentration of AgNPs inhibited N_2_O reduction by *P. denitrificans* R-1.

**Fig. 7.**
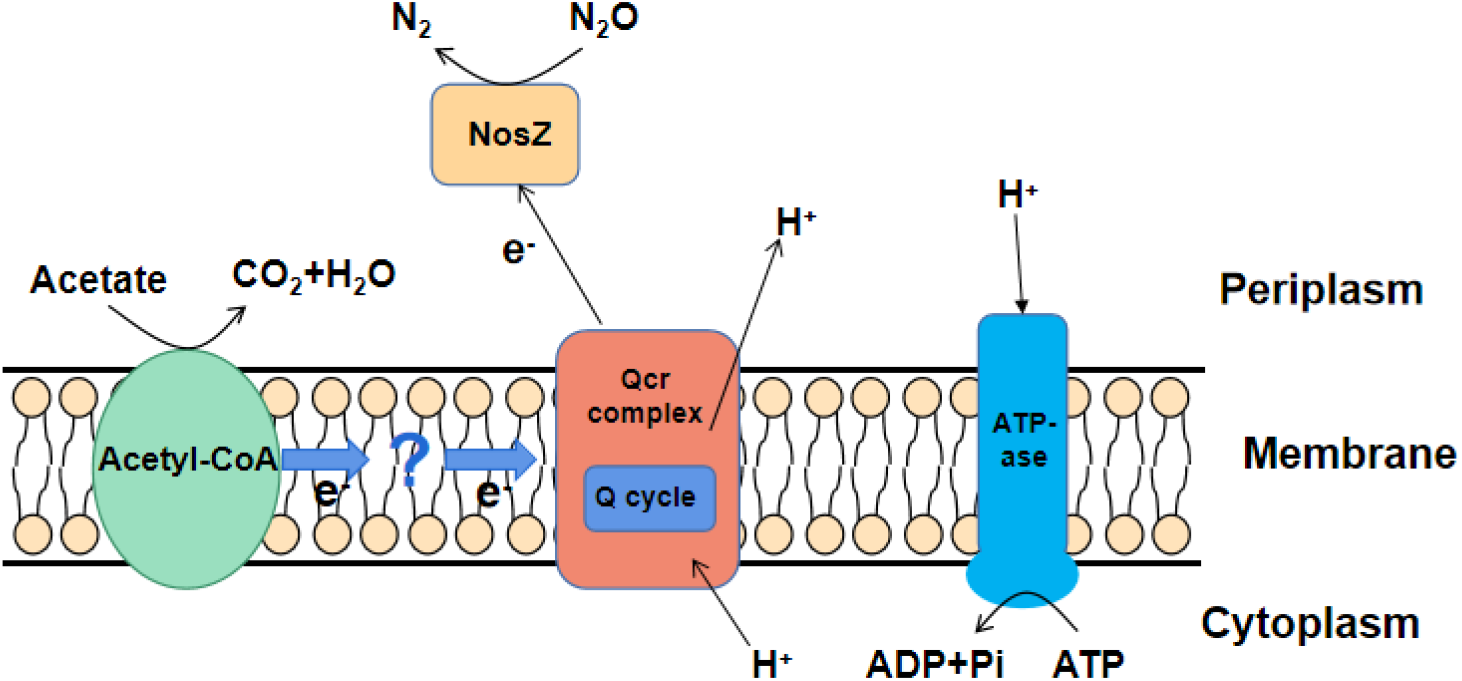
*P. denitrificans* R-1 N_2_O respiratory electron transport model using sodium acetate as electron donor. For simplicity, only major enzymes are shown. Electron transfer chain also includes nosD, Y, L, X, and R which are not shown, and unknown enzymes cannot be ruled out.

## DISCUSSION

### Oxidation of electron donor coupled to N_2_O reduction by *P. denitrificans* R-1

Microbial N_2_O respiration is an important process in the N cycle. Many microorganisms respond to N_2_O as an electron acceptor (8, 9, 11, 22). In the current study, we found that *P. denitrificans* R-1 could reduce N_2_O via the oxidation of electron donors. Sodium acetate, ethanol, sodium propionate, sodium pyruvate, sodium lactate, sodium succinate, and glucose acted as effective electron donors to support N_2_O reduction by *P. denitrificans* R-1, and sodium acetate was the strongest electron donor for N_2_O reduction. According to previous studies (23, 24), acetate can be converted into acetyl-CoA and then directly integrated into the TCA cycle for degradation in denitrifying bacterial cells; therefore, the utilization rate of acetate is more efficient than that of other electron donors, resulting in a higher N_2_O reduction rate. We also studied the coupling relationship between the oxidation of the electron donors, sodium acetate or sodium lactate, and the reduction of the electron acceptor, N_2_O. Linear fitting analysis also confirmed that the oxidation of the electron donor and the reduction of the electron acceptor showed a typical coupling relationship (R^2^ > 0.9). The redox potentials of various electron donors are related to the different N_2_O reduction efficiencies.

When sodium acetate and sodium lactate are used as electron donors, the reduction process of N_2_O conforms to the chemical equation in Table 1. Theoretically, 1 mol of sodium acetate can provide 8 mol of electrons and support 4 mol of N_2_O reduction. 1 mol of sodium lactate can provide 12 mol of electrons, supporting 6 mol N_2_O reduction. The ratios of the acetic acid and sodium lactate consumed by *P. denitrificans* R-1 to the amount of N_2_O reduced were 1:0.89 and 1:2.07. The bioavailability efficiency of sodium acetate and sodium lactate reached 22.25% and 34.50%, respectively. This result indicates that the oxidation of acetate or lactate was sufficient for energy conservation when N_2_O was completely reduced to N_2_.

**Table 1.** Theoretical and actual ratios of electron donor to acceptor in the Oxidation-Reduction Reaction.

| Electron donor | Electron acceptor | Oxidation-Reduction Reaction | Ratio of donor to acceptor (Mole/Mole) |  |  |
| --- | --- | --- | --- | --- | --- |
|  |  |  | Theoretical | Experimental | efficiency |
| Acetate | Nitrous oxide | $\frac{1}{8}CH_3COO^- + \frac{1}{2}N_2O \rightarrow \frac{1}{8}CO_2 + \frac{1}{8}HCO_3 + \frac{1}{2}N_2 + \frac{3}{8}H_2O$ | 1:4 | 1:0.89 | 22.25% |
| Lactate | | $\frac{1}{12}CH_3CHOHCOO^- + \frac{1}{2}N_2O \rightarrow \frac{1}{6}CO_2 + \frac{1}{12}HCO_3 + \frac{1}{2}N_2 + \frac{1}{6}H_2O$ | 1:6 | 1:2.07 | 34.50% |

### Energy conservation from dissimilatory N_2_O reduction by *P. denitrificans* R-1

Cell growth depends on the supply of energy and nutrients. In contrast to previous reports (2, 11, 25), the growth of *P. denitrificans* R-1 was observed only when N_2_O was reduced as the sole electron acceptor. The OD_600_ value increased from 0.0675 to 0.1925 within 24 h of incubation (Fig. 2), suggesting that the coupled oxidation of acetate to N_2_O reduction provided energy for the growth and metabolism of *P. denitrificans* R-1. A previous study showed that some denitrifiers can grow by N_2_O reduction using N_2_O as the sole electron acceptor, known as N_2_O respiration bacteria (NRBs). Most of these bacteria are facultative anaerobes and harbor a clade II N_2_O reductase, including *Wolinella succinogenes* (26), *Campylobacter fetus* (27), *Anaeromyxobacter dehalogenans* (6)*, Bacillus vireti* (8), *Dechloromonas aromatica* (28), *Dechloromonas denitrificans* (28), and *Azospira* sp. strain I13 (10). However, N_2_O respiration is widely underexplored, and only a few NRBs of traditional denitrifiers with clade I N_2_O reductase have been shown to grow through N_2_O reduction. Some NRBs possess *nrfA*, a key functional gene for dissimilatory nitrate reduction to ammonium (DNRA), suggesting that N_2_O reduction is coupled with nitrogen fixation, in which N_2_O is first reduced to N_2_, and then N_2_ is further reduced to ammonium nitrogen and integrated into the cell biomass (29). In addition, Park et al. showed that *G. auruantiaca* T-27 was able to reduce N_2_O when O_2_ was depleted and O_2_ was initially present, but no growth was observed (11). A plausible explanation for this lack of growth is that obligate aerobic microorganisms with *nosZ* may utilize N_2_O as a temporary surrogate for O_2_ to survive periodic anoxia. In the present study, our results suggest that *P. denitrificans* R-1, a traditional denitrifier with clade I N_2_O reductase, can grow when N_2_O is reduced coupled with electron oxidation; therefore, *P. denitrificans* R-1 can be called a NRB.

### Electron transportation system for microbial N_2_O reduction

Respiratory inhibitor experiments showed that the electron transfer of *P. denitrificans* R-1 in the N_2_O reduction process did not involve the conventional respiratory electron transfer enzyme complexes I and II (Fig. 3). Previous studies have reported that the electron transport chain of classic *nosZ*-Ⅰ type of denitrifying bacteria is located on the cell membrane, including QCR compounds, the Q circulation system, cytochrome C, nosZ reductase, and *nos* genes encoding proteins (NosR, - X - C, - D, F, -Y, and -L) (2). Genomic data showed that *P. denitrificans* R-1 has a similar *nos* gene cluster (NosR, -Z, -D, -F, -Y, and -L) (Fig. 4), suggesting that *P. denitrificans* R-1 has electron-transfer protein components similar to those of classic NosZ-I-type denitrifying bacteria.

It has been shown that three proteins, NosD, NosY, and NosF, encoded by *nos* gene clusters, may constitute a complex transporter that binds to the cell membrane and couples ATP hydrolysis; however, it is unclear whether they can play the role of transporters (2, 17, 30). NosL is a Cu-containing outer membrane lipoprotein that is closely related to the NosDYF complex. Studies have suggested that NosL may provide Cu for NosZ (17). NosR also participates in electron transfer. FeS and FMN are distributed at both ends of this protein and can transfer low-potential electrons from the cytoplasm across membranes to NosZ located in the pericytoplasm, which is an electron transfer pathway independent of Qcr (2). NosX is a signal peptide containing a Tat sequence that exists in the periplasmic space and contains a flavin protein with FAD as a co-group. NosX is mainly involved in the biogenesis of NosR and is a cofactor of the FMN terminal of NosR. NosX is associated with the influence of ApbE proteins in Fe-S centers, and ApbE has been shown to be a flavin donor in NosR (2, 31).

Based on inhibitor experiments and genomic analysis, we deduced an electron transfer model for the N_2_O reduction growth of *P. denitrificans* R-1 (Fig. 7). Electrons produced by the conversion of acetic acid are transferred through the electron transport chain, generating an electrochemical force on the membrane and driving ATP synthesis. However, the exact mechanisms underlying microbial N_2_O respiration remain unclear.

### Potential ecological significance of microbial N_2_O reduction

N_2_O is one of the most significant forms of nitrogen pollution. It is currently the third largest adult greenhouse gas emitted and the largest man-made source of stratospheric ozone (32). Many biological and abiotic processes contribute to the production of N_2_O in the biosphere. However, the consumption of N_2_O in the environment has been largely attributed to denitrifying microorganisms. *nosZ* encoding the N_2_O reductase enzyme NosZ, responsible for N_2_O reduction to dinitrogen, are now known to include two distinct groups: the well-studied clade I, which is a typical denitrifier, and the novel clade II, possessed by diverse groups of microorganisms, most of which are non-denitrifiers (6).

Denitrification pathways are highly modular. Reduction of N_2_O by typical denitrifying bacteria occur after the production of N_2_O. Moreover, the reduction of N_2_O is an independent process (33). Our study showed that the *nosZ* type I bacterium, *P. denitrificans* R-1, can respire N_2_O as the sole electron donor. Thus, the modular N_2_O reduction process of clade I denitrifiers not only can consume N_2_O produced by themselves but can also consume the external N_2_O generated from non-denitrification biological or abiotic pathways under suitable conditions, which is critical for controlling the release of N_2_O from ecosystems into the atmosphere.

## ACKNOWLEDGMENTS

This work is funded by the National Natural Science Foundation of China (No. 42276130 and No. 42006122), Natural Science Foundation of Guangdong Province (No. 2021A1515011548, No. 2022A1515010539, and No. 2023A1515012424), Basic and Applied Basic Research Foundation of Guangdong Province (No. 2020A1515110597), the Innovation Team Project of Guangdong Provincial Department of Education (No. 2021KCXTD016), and the Basic Research Project of Guangzhou City and School Joint Funding (No. 202201020579, No. 202201020542 and No. 202201020580).

